# Climate-Driven Reshuffling of Butterfly Communities: Body Size Declines and Community Homogenization in a Rapidly Warming Switzerland

**DOI:** 10.1101/2025.02.06.636896

**Authors:** Konstantina Zografou, Vassiliki Kati, Eva Knop

## Abstract

Climate change has been reshaping natural communities by driving species to change their ranges in response to warming temperatures. In Switzerland, the rate of warming since 1950 has approached nearly double the global average, making these impacts important for community composition and ecosystem function. Using the Community Weighted Length Index (CWLI) as the body size estimator, previously extracted by museum specimens, we studied two decades of butterfly community data that have been inventoried by the Biodiversity Monitoring Switzerland program. We checked for deviations from the expectation of random trends in the CWLI using null models. We modeled the long-term patterns at each site and the overall long-term patterns through linear regression and mixed-effects models alongside the model of the relationship between wing length, elevation, and abundance. CWLI of Swiss butterfly assemblages dropped by 5.2% over two decades, suggesting that warmer temperatures are differentially affecting large-bodied species. While CWLI trends were significantly declining at most sites (73%), a subset of sites (27%) showed trends of increasing CWLI, sometimes associated with cooler microclimates either in forests or at higher elevation. Our results further indicate a progressive homogenization of butterfly communities, marked by smoother CWLI slopes and reduced CWLI range over time. These results underline complex relations of climate-driven range shifts with body-size dynamics and community restructuring, emphasizing the importance of understanding the processes to mitigate biodiversity loss in thermally sensitive ecosystems.

## Introduction

Climate-induced changes in the environment are pushing species around the world to adjust their geographical range limits to match their optimum thermal preferences (Parmesan 2006, Chen et al. 2011, Scheffers et al. 2016). Distributional changes have been widely documented in various plant and animal taxa and it is expected that species that are unable to adapt or move will go locally or regionally extinct (Maggini et al. 2011). In addition, the speed and timing of responses are found to greatly vary across space and taxonomic group, resulting in the disruption of important ecological interactions, and the formation of new assemblages of species may take place (Pecl et al. 2017). Evidence shows that the rate of range shifts increases with increased warming (Chen et al. 2011) and is expected to continue increasing as global climate change becomes more pronounced (Urban 2015).

Interspecific range shifts are likely to give rise to newly organized biological communities with the co-occurrence of old species in new mixtures (Walther 2010). The examination of such new communities generally involves testing temporal or spatial variation in morphological, physiological, or behavioral traits along environmental gradients such as elevation or latitude. For example, in Borneo, climate-associated range shifts were found to decrease the average body size of novel moth communities, such that smaller species became more abundant and large-bodied associated with colder climates (consistent with Bergmann’s rule) declined (Wu et al. 2019). In a similar application, readings utilizing parameters like the species temperature index (STI) established changes in the composition of the community also at temporal scales, where the warm-adapted species gradually replaced populations of birds and butterflies in cold-adapted species over time (Devictor et al. 2012, Zografou et al. 2014, Bonelli et al. 2022). The speed of the change varies, however, depending on the group studied in the community and the sensitivity of the study area. High-altitude or otherwise thermally more extreme localities are particularly at risk for this effect (Habel et al. 2023).

Trophic interactions can depend importantly on changes in body size, and body size also underlies shifts in dietary requirements, predator-to-prey body size ratios (Yodzis and Innes 1992, Gibert and DeLong 2014), and ecological roles more generally. Changes in body size could further result in changes in the flow of biomass and the stability of food webs (Emmerson and Raffaelli 2004); however, empirical data regarding the impact of climate-driven range shifts on community body size is surprisingly scarce. This applies in particular to insects, which comprise most of the terrestrial biodiversity and are key to energy transfer within ecosystems (Prather et al. 2013). The assessment of these impacts is of uttermost importance, especially in areas where climate warming sensitivity is much higher than the global average. Warming in Switzerland has occurred at about twice the global average rate since 1950 (Foen 2020). This trend makes the country an ideal location to explore the effects of climate change on living populations. Rapid warming in Switzerland has begun to alter conditions -ecological and human alike-in increasingly perceivable ways (Vittoz et al. 2013).

Terrestrial ectotherms, particularly butterflies, have already demonstrated significant responses to climate change in Switzerland. These include advancing their phenology (Altermatt 2010), colonizing regions with warmer climates (Boillat 1992, Juillerat 2005), and shifting their distributions to higher elevations (Hohl 2006, Pasche et al. 2007). However, evidence for community-level body size reduction in response to rising temperatures is still lacking. Despite mixed evidence of trends in body size for butterflies, with reports varying from increases (Wilson et al. 2023) to decreases (Bowden et al. 2015, Wu et al. 2019, Semsar-kazerouni et al. 2022), there is strong evidence of community reshuffling in the sense that large size butterflies will fly out and smaller size butterflies will fly in (but see (Merckx et al. 2018) for an inverse pattern for butterflies along urbanization gradients). In that sense, we would expect to find a decreasing community size trend over time.

Here, we sought to provide empirical evidence on community reshuffling by addressing directional shifts in community body size over the years. We used butterfly surveys from the Biodiversity Monitoring Switzerland program (BDM) and butterfly forewing length from museum specimens (unpub. data Zografou), to explore changes in community composition due to interspecific shifts in range. This study unravels the web of interactions among climate-driven range shift, body size change, and their consequences for community reshuffling in a region that is rapidly warming.

## Methods

### Butterfly data

The study was based on data gathered within the Swiss Biodiversity Monitoring (www.biodiversitymonitoring.ch). We limited the sample selection for our analyses to records of day-flying butterflies. The analyses considered 177 of the 192 species that had been recorded, including the day-flying species of the family Zygaenidae. The temporal coverage of 20 years of data, from 2003 to 2023 inclusive, was gathered within standardized butterfly surveys along transects within 450 systematically arranged sites of 1 km^2^ throughout Switzerland. Each of these was surveyed at 5-year intervals, 5 to 7 times per year. Because the surveys required are fewer for higher elevations with shorter flight seasons, the number of surveys per year varied with the elevation.

### Wing length dataset

We used measurements of wing length for specimens removed from museum specimens housed in two major butterfly collections: the Entomological Collection of the Swiss Federal Institute of Technology located in Zürich (ETHZ) and the Goulandris Natural History Museum (GNHM). Initially, morphometric measurements were carried out manually and then with the help of computer vision for automation, which provided a data set that includes morphometric data of 31,920 unique specimens representing 588 species, collected in 88 countries between 1840 and 2010 (unpub. data Zografou). Forewing length serves as a reliable proxy for body size in museum specimens (Brehm et al. 2019, Minter et al. 2024), and body size reduction is recognized as the third universal biological response to climate change (Sheridan and Bickford 2011).

To set our analysis in the context of Switzerland, we then filtered the dataset to include only specimens collected within the country, so that we have estimated mean forewing lengths for 164 of the 177 butterfly species. Mean lengths for two additional species (*Argynnis pandora, Boloria selene*) were derived from specimens collected outside Switzerland. Based on this, we systematically excluded those 17 distinct species of diurnal moths, together with 13 butterfly species for which data on length were missing (namely, *Cacyreus marshalli, Erebia albergana, Erebia arvernensis, Erebia flavofasciata, Erebia pronoe, Erebia sudetica, Erebia triarius, Euphydryas intermedia, Hamearis lucina, Melitaea asteria, Melitaea deione*, and *Parnassius phoebus*). This reduced, high-quality dataset was hence taken and formed the basis of our further statistical analysis.

### Community Weighted Length Index (CWLI)

The community-weighted length index (CWLI) was calculated for each site and year, considering only species with available length measurements (see above). The weighted mean was used, with weights corresponding to the number of individuals for each species using the following formula:

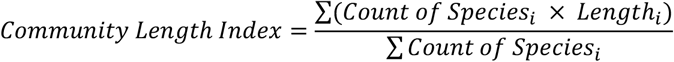

where *Count of Species*_*i*_ is the abundance of species *i* in the community and *Length*_*i*_ is the average length of species *i*.

### Statistical approaches

Null models were used to create randomizations that allow for comparison with observed patterns. The goal was to see whether the observed changes in Community Weighted Length Index (CWLI) can be explained by random processes or are likely due to other ecological factors such as climate change or land use change. To secure a distribution of expected index values under random conditions, we shuffled species abundances across the sites and years (999 randomizations), while maintaining the total abundance and species richness per site-year to break any temporal or spatial correlations.

We used linear regressions to assess long term trends at each site. In addition, a mixed effects model with CWLI as the response variable, and year, elevation and their interaction term as fixed effects, and sites as a random intercept to account for site-specific variations was fitted for assessing an overall trend across all sites.

To disentangle community reshuffling trends, we modeled abundance data to wing length and elevation. We first used a generalized linear model with a poisson distribution for count data (REF). However, model diagnostics detected overdispersion and multicollinearity. To improve the fit of the model and overcome dispersion and collinearity issues we applied a negative binomial distribution after having centered and scaled the variables. We run diagnostic plots to ensure that assumptions of linearity, homoscedasticity, and normality were met.

All statistical analyses were performed through R version 4.3.2 (R Core Team 2024). The R packages dplyr (Wickham et al. 2023), ggplot2 (Wickham 2016), broom (Robinson D et al. 2024), tidyverse (Wickham et al. 2019), nlme (Pinheiro et al. 2021), tidyr (Wickham H et al. 2024), mass (Venables and Ripley 2002), and gridExtra (Baptiste 2015) were used for data handling, data analysis, and plotting.

## Results

Null models suggested that observed CWLI significantly deviates from the null model distribution in 24 occasions (Table S1) and that these occasions are visualized to be closer to the higher CWLI values (Fig. 1).

**Figure 1.**
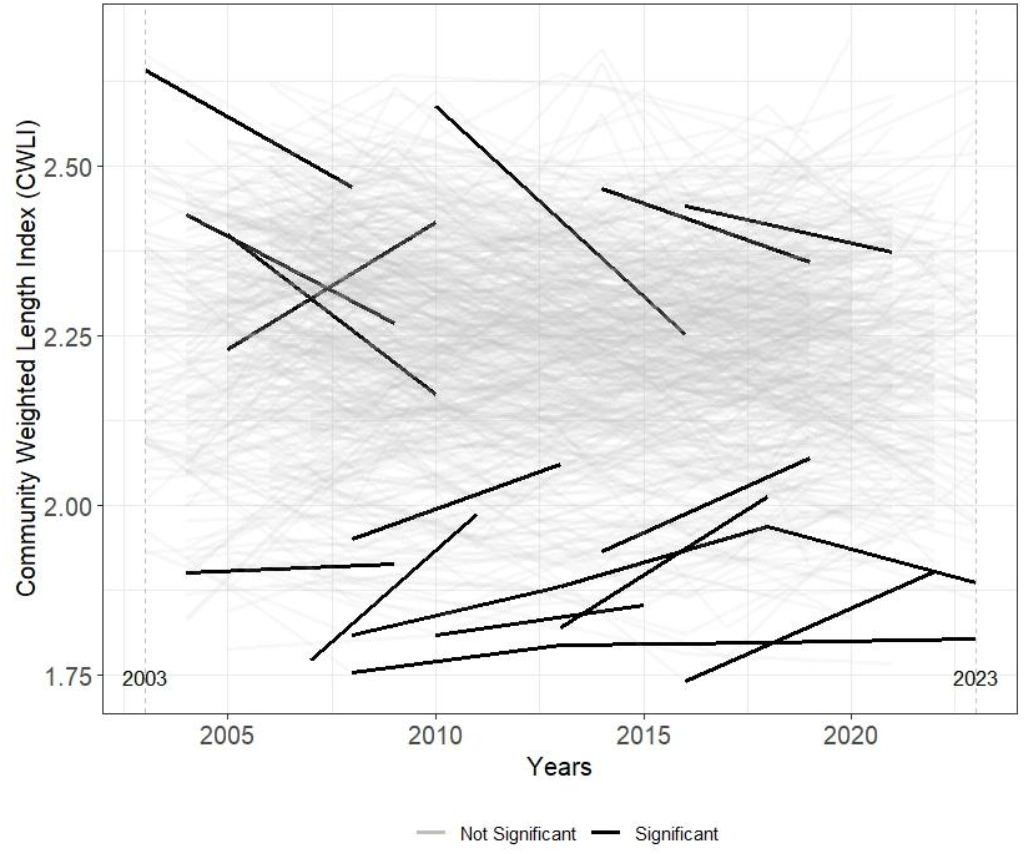
The plot overlays the observed CWLI with the null mean for each site (450 in total) over the studied period (2002 – 2023). Sites and years with *P*-values *< 0.05* are flagged with black color showing significant differences (two tailed test). Dashed vertical lines in 2003 and 2023 mark the start and end years of the survey period.

We found a small number of sites (8.2%) to have a significant trend when testing for shifts in CWLI (Table S2). However, it becomes clear, when focusing on significant trends, that decreasing trends (73%) favored over the years, while 27% of site-specific CWLI only showed a positive trend (Fig. 2).

**Figure 2.**
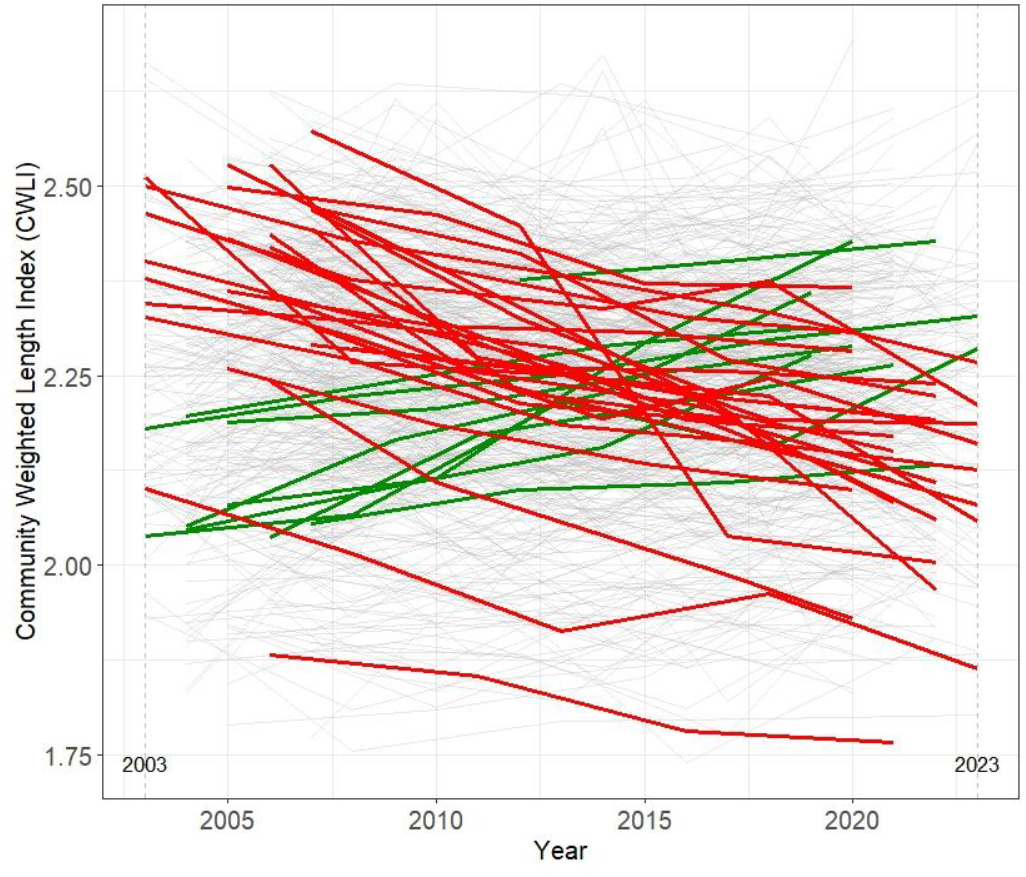
Trends in Community Weighted Length Index (CWLI) over time across survey sites. Gray lines represent sites with no trend, green lines indicate sites with an increasing trend, and red lines indicate sites with a decreasing trend. Dashed vertical lines at 2003 and 2023 mark the start and end years of the survey period.

All three relationships between CWLI, elevation, year and their interaction term were significant (Table S3). Using performance package (Lüdecke et al. 2021) we estimated the conditional variance explained by both fixed and random effects (R^2^ = 0.645) and the marginal variance explained by fixed effects (R^2^ = 0.232). CWLI was found to decrease over the years (β_years_ = -0.005, *P* < 0.001) (Fig. 3) and, surprisingly, was also found to decrease with elevation (β_elevation_ = -0.006, *P* < 0.001) (Fig. 4). The dynamic interaction between temporal trends and altitudinal gradients is captured in Figure 5 as a dynamic alignment (convergence) of CWLI over the years and in Figure 6 as a progressive smoothness on slopes, starting with the sharper slope in 2003 and ending to the smoother slope in 2023.

**Figure 3.**
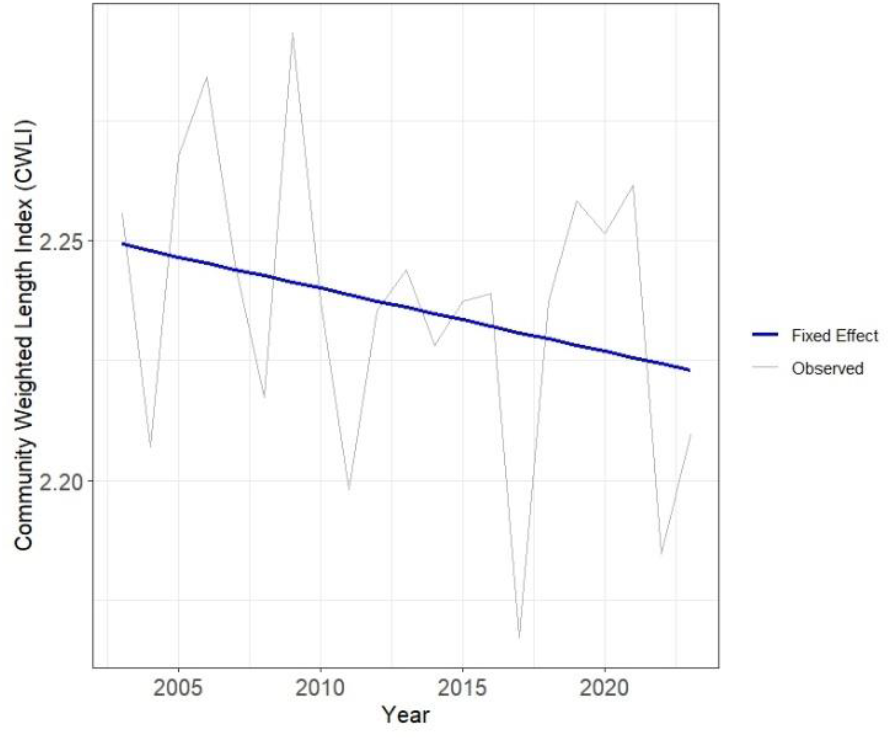
General trend in Community Weighted Length Index (CWLI) over time. Gray line represents the observed trend for CWLI data considering all sites and the blue line represents the predicted trend based on the fixed effects of the linear mixed-effects model.

**Figure 4.**
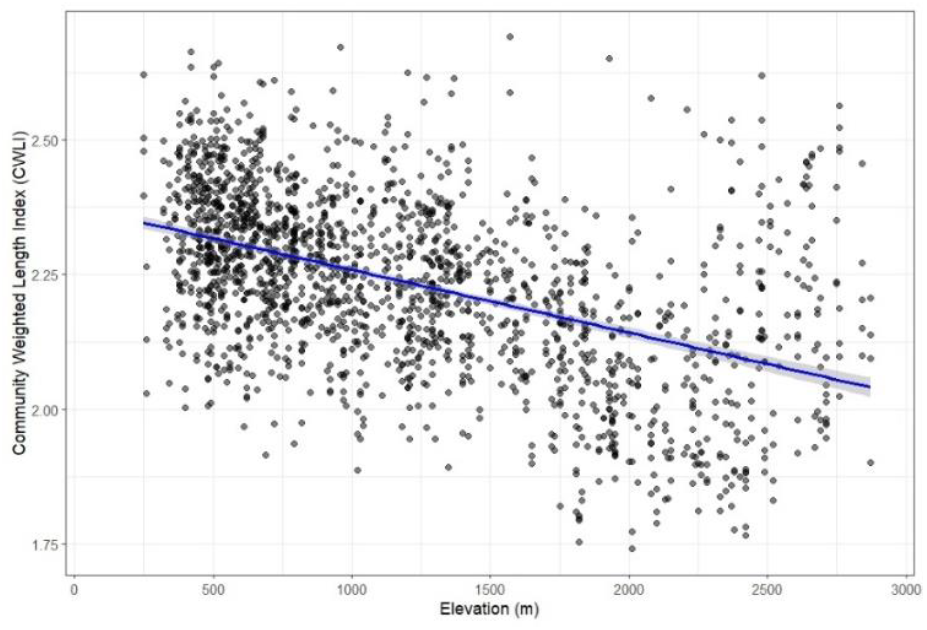
The effect of elevation to community weighted length index (CWLI). The scatter plot depicts the observed CWLI values (grey points) across a gradient of elevations (in meters). The blue line represents the fitted linear regression model, indicating the overall trend in CWLI with increasing elevation.

**Figure 5.**
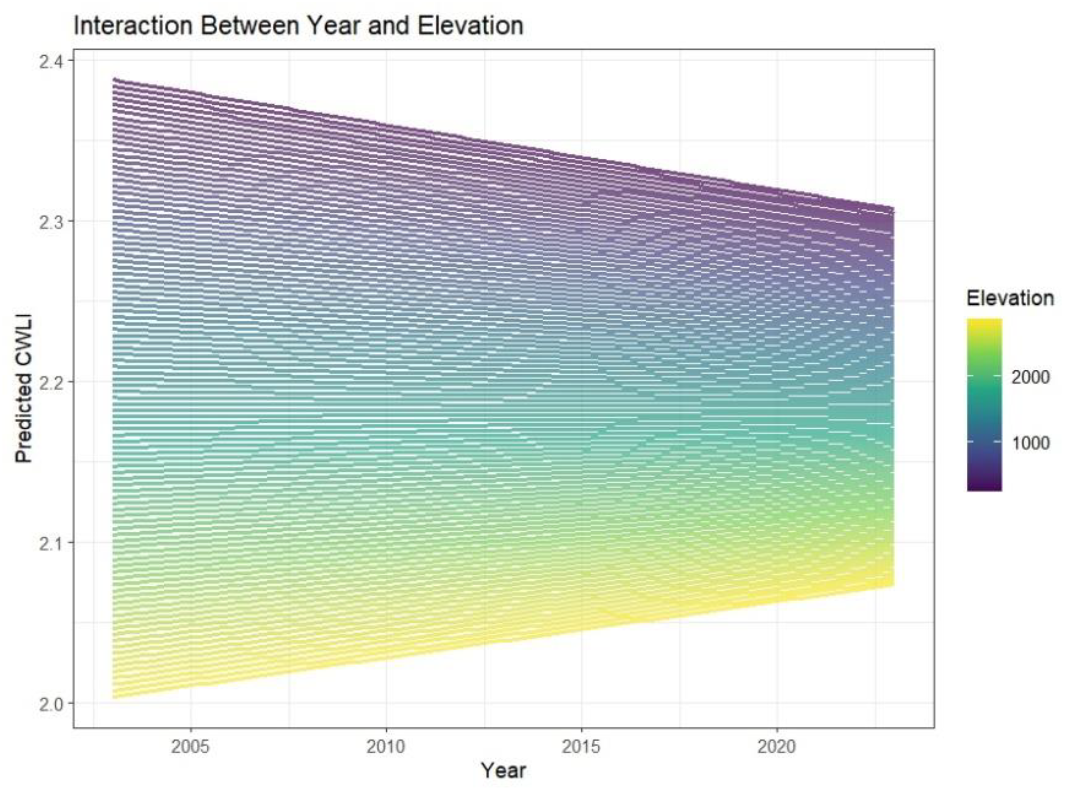
Interaction between year and elevation predicting the Community Weighted Length Index (CWLI). The plot illustrates how the predicted CWLI changes over time across different elevation gradients. Each line corresponds to a specific elevation, highlighting the variability in CWLI trends across altitudinal gradients. Warmer colors (yellow) indicate higher elevations, while cooler colors (purple) represent lower elevations.

**Figure 6.**
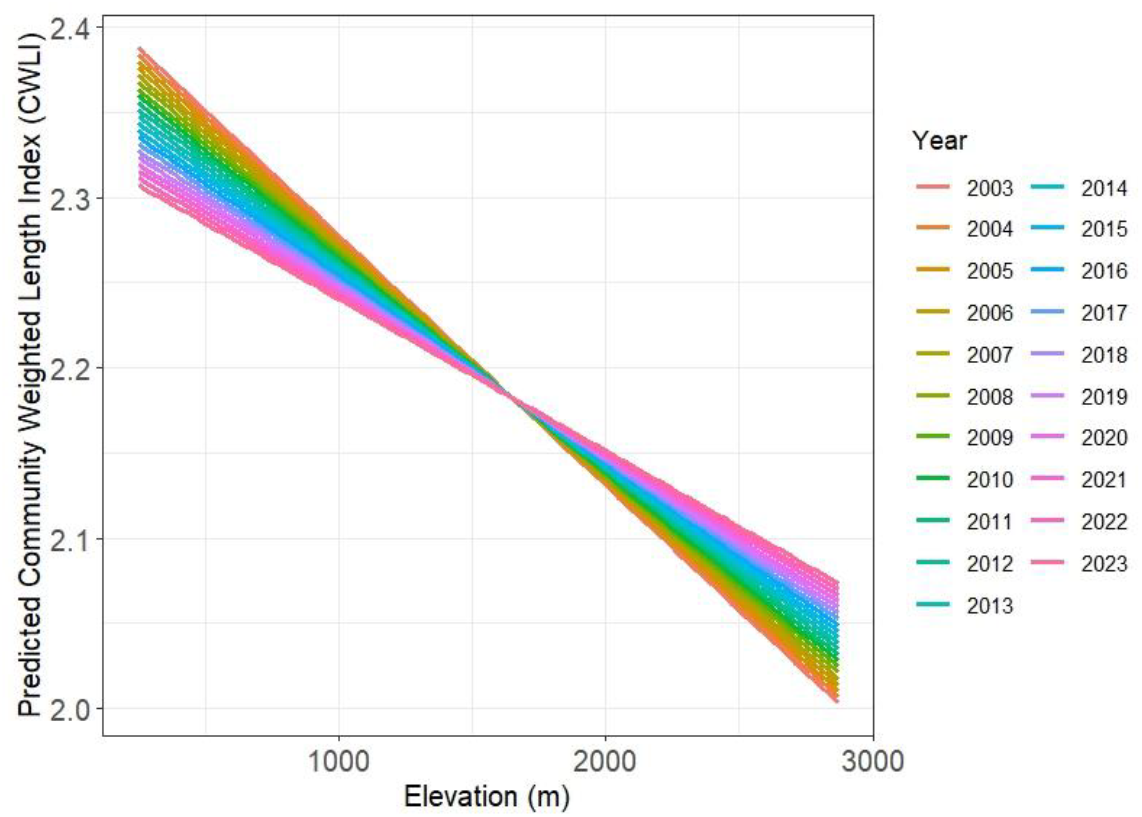
Effect of elevation on the predicted Community Weighted Length Index (CWLI) across years. The plot shows predicted CWLI values based on the interaction between elevation and year, derived from the linear mixed-effects model. Each line represents the predicted CWLI for a specific year, as indicated by the color legend. The model accounts for random effects associated with site-specific variation and shows a progressive smoothness on slopes, starting with the sharper slope in 2003 and ending to the smoother slope in 2023 across elevation.

We found abundance to be driven by both elevation (β_elevation_ = 0.11, *P* < 0.001) and wing length (β_wing length_ = -0.06, *P* < 0.001). The most outstanding output, however, is their interaction effect (β_elevation_ = - 0.07, *P* < 0.001), where larger species lower their abundance while moving up to higher altitudes justifying the decreasing CWLI we found along elevation (Table S4, Fig. 7). Model fitted well (x^2^ = 61758, *P* = 1), with no evidence of overdispersion (dispersion ration = 0.9) and/or multicollinearity.

**Figure 7.**
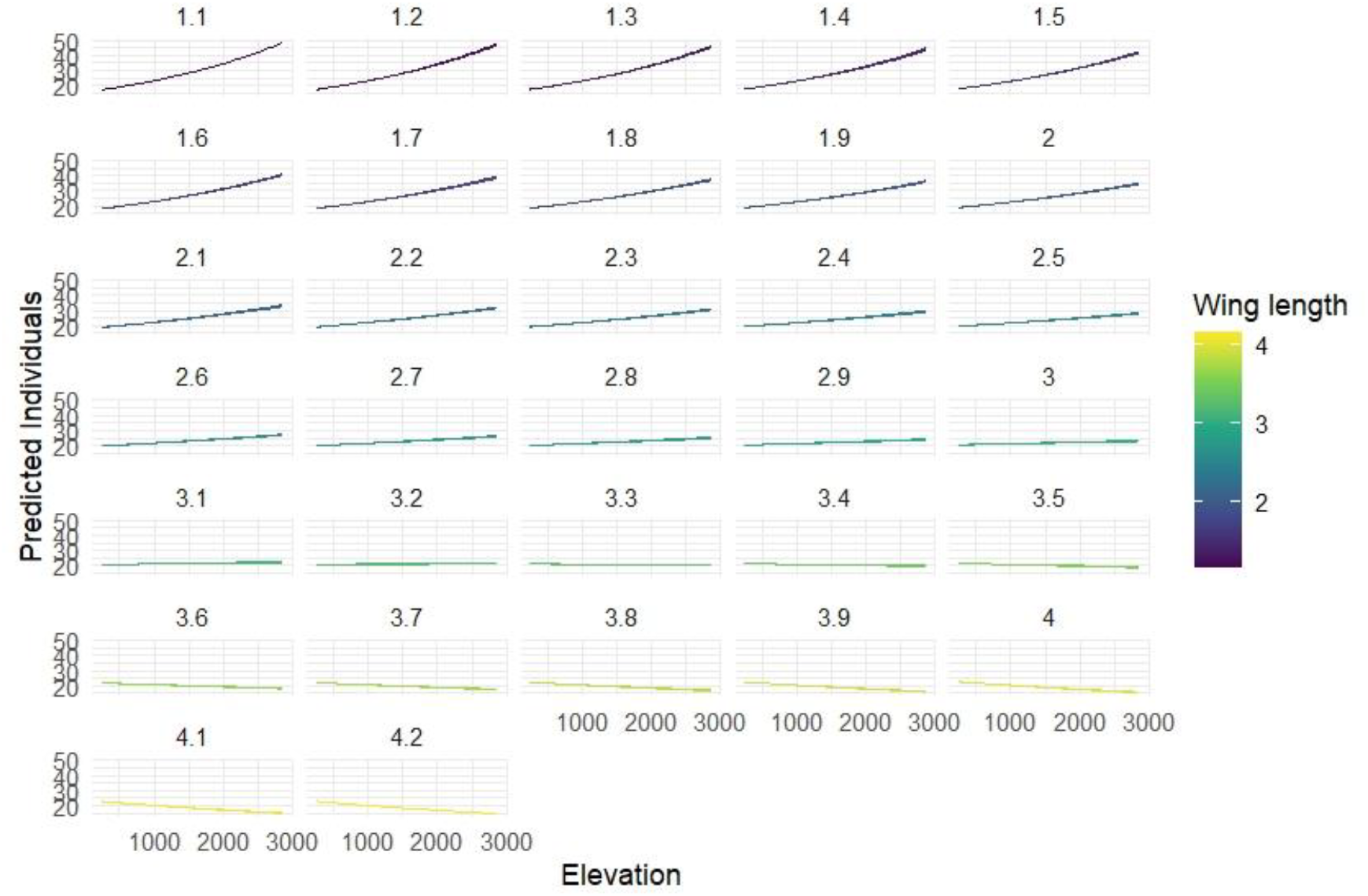
The predicted number of butterfly individuals across a gradient of elevation and wing length (forewing length, used as a proxy for body size). The predictions were generated using a generalized linear model (GLM) that incorporates elevation, wing length, and their interactions as explanatory variables.

## Discussion

We observed a clear temporal decline in the Community Weighted Length Index (CWLI) of butterfly communities across Switzerland during the last two decades. Results underpin the dynamic interplay of temporal trends with altitudinal gradients, driving the homogenization of butterfly communities in Switzerland. The progressive reduction of CWLI values across the elevations coupled with the contraction of the CWLI range over time are indicative of crucial ecological shifts such as community reshuffling and loss of distinctiveness through interactions between smaller and larger species under changing climate.

Overall, the CWLI of Swiss butterfly assemblages dropped by 5.2% over two decades. At the site level, a low percentage (8.2%) of sites had strong evidence for significant trends. Of these, most sites (73%) had a declining trend consistent with the overall trend, but an important minority (27%) had increasing trends, suggesting localized deviations from the general trend of decline. This reduction in CWLI is according to the size-temperature rule (Atkinson 1994, Verberk et al. 2021) and is most directly attributable to rising temperatures. Switzerland has warmed noticeably in the last couple of decades, and its average temperatures have increased at almost twice the global rate (Foen 2020). More intense warming should therefore enhance the decline in size at the community level.

Interestingly, the observed increases in CWLI at some sites reflect a gradual replacement of smaller species by larger ones. Spatial inspection of these locations indicated that six were situated within forested regions, three in elevated pastures exceeding 1500 meters, and one in agricultural land. These trends are most likely the product of microhabitat buffering through forest shaded and cooler habitats that provide refuge from drought and extreme heat (Bruschini et al. 2024). Open lowlands are highly exposed to these stressors, and this amplifies ecological pressures on butterfly populations.

The findings of this study introduce two key insights into the temporal decrease of the Community Weighted Length Index (CWLI) in butterfly assemblages along elevational gradients. First, the complex relationship between altitudinal gradients and temporal tendencies reveals a smooth transition towards more stable CWLI slopes over the course of the study period (Fig. 6). This trend is accompanied by a uniform reduction in the range of CWLI values over time (Fig. 5). Second, the altitude decrease of CWLI appears primarily as the driving force of a progressive decrease in the proportion of larger species at increased altitudes (Fig. 7).

The observed decrease in variability of CWLI and decreasing range of values over the years are signs of continued homogenization of Swiss butterfly assemblages. Similar trends have been noted in the Italian Alps but have been attributed to changes in other ecological characteristics, such as mobility and thermal tolerance (Bonelli et al. 2022). Less distinctive communities may result from intraspecific changes to smaller body sizes, which could serve as a mechanism to meet elevated energetic demands due to increasing temperature (Atkinson 1994, Sheridan and Bickford 2011). Community reshuffling can also play an important role, wherein small species migrate to greater altitudes, and the abundance or range of large species is reduced or they go extinct due to warming climate constraints (Bergmann 1847, Forster et al. 2012).

Based on species ecothermal rules (Bergmann 1847, Atkinson 1994), we expect to find smaller species at lower elevations and/or latitudes and larger species at higher elevations and/or latitudes. But, the increasingly driven by human-mediated climate change is already forcing organisms to shift their range upward towards favorable conditions (Illán et al. 2012) driving communities to a gradual homogenization. Especially in mountainous regions like Switzerland, warming effect is stronger and species communities often show particularly pronounced responses and movements (Chen et al. 2011). For instance, (Habel et al. 2023) showed that lowland butterflies have shifted their average occurrence by more than 300 m uphill in northern Alps, while (Freeman et al. 2018) confirmed the upslope shift not only for ectotherms but also endotherms and vascular plants. Moreover, Freeman et al. (2018) found mountaintop species to be strongly affected by the rising temperatures as elevational ranges and available area significantly decreased for those organisms.

Our results support the community reshuffling with smaller species to increase their proportions at the expense of larger species along the elevation (Fig. 7). The influx of species from lower elevations likely intensifies competition for available larval and adult food resources which is stressing for larger species, often associated with increased food consumption due to their higher metabolic demands (Calder 1996, Speakman 2005). It is also plausible that some species have already reached the highest elevations within their range, leaving no further room for upward migration, considering also the decrease of available areas at the mountaintops (Parmesan and Yohe 2003, Freeman et al. 2018). Or simply larger butterfly species may shift their distributions toward higher latitudes, gradually disappearing from the study area. Alternatively, larger species might abandon open areas above treelines, such as mountaintops, and relocate to cooler, shadier environments, including woodlands or areas in close proximity to water sources (e.g., lakes, rivers) as a response to rising temperatures (Pateman et al. 2016). Butterflies have been found to change their habitat preference as per annual climatic fluctuations, where they prefer cooler, shaded habitats like forests in hot years and shift towards warmer, open habitats in cold years (Suggitt et al. 2012). Such habitat shifts may be able to explain the observed rise in Community Weighted Length Index (CWLI) we found.

Since intraspecific variation in body size was not accounted for, it is likely that even larger differences would emerge when examining individual species categories in the context of community length reduction. The patterns, however, uncovered here suggest that changes in community size could be driven both by direct thermal stress and resource-related constraints and point to the complex interplay of factors underlying community-level changes. We emphasize the importance of directional conservation initiatives to counter the effects of climate change on biological diversity and ecological stability in mountain ecosystems, which are extremely vulnerable to increasing temperatures.

## Code availability

The R-script of the analysis is available at Zenodo (DOI: 10.5281/zenodo.14772406).

## Acknowledgements

The Swiss Federal Office for the Environment (FOEN) kindly provided the BDM data. We thank the dedicated team who conducted the fieldwork for the Swiss Biodiversity monitoring program.

## Author contributions

K.Z. conceived and designed the study. E.K. facilitated the use of BDM data, K.Z. performed statistical analyses and prepared visualizations. K.Z. secured funding and V.K. supervised the project. The manuscript was originally drafted by K.Z. and V.K. and E.K. contributed to the interpretation of results, manuscript revisions, and approved the final version for submission.

