## Supporting Information for "Climate-Driven Reshuffling of Butterfly Communities: Body Size Declines and Community Homogenization in a Rapidly Warming Switzerland"

This file includes supplementary tables, and figures (diagnostic plots) that support the conclusions of the main manuscript. It is intended to provide transparency and reproducibility for the reported findings.

**Contents:**

Table S1 – S4

Figure S1- S3

For any questions regarding this material, please contact []

**Table S1**. Significant changes between observed CWLI (Community Weighted Length Index) and the null model using a z-score and p-value. A z-score shows how many standard deviations the observed CWLI is from the null mean and p-values < 0.05 significantly deviates from the null model distribution.

| id_site | year | CLI | total_count | null_mean | null_sd | z_score | *p-value* |
| --- | --- | --- | --- | --- | --- | --- | --- |
| 593086 | 2005 | 2.23 | 140 | 1.95 | 0.139 | 2.01 | 0.0444 |
| 605230 | 2003 | 2.64 | 378 | 2.24 | 0.181 | 2.21 | 0.027 |
| 623126 | 2010 | 2.59 | 933 | 2.16 | 0.205 | 2.08 | 0.0373 |
| 623126 | 2020 | 2.69 | 903 | 2.22 | 0.202 | 2.34 | 0.0195 |
| 629118 | 2004 | 2.43 | 1112 | 2.16 | 0.104 | 2.55 | 0.0109 |
| 629118 | 2014 | 2.47 | 3327 | 2.18 | 0.12 | 2.39 | 0.0169 |
| 653118 | 2021 | 1.87 | 3644 | 2.13 | 0.116 | -2.29 | 0.0218 |
| 671158 | 2010 | 1.81 | 1409 | 2.1 | 0.115 | -2.58 | 0.01 |
| 677182 | 2005 | 2.4 | 913 | 2.17 | 0.107 | 2.16 | 0.0305 |
| 689142 | 2016 | 2.44 | 1498 | 2.19 | 0.125 | 2 | 0.0451 |
| 689166 | 2008 | 1.75 | 1119 | 2.05 | 0.137 | -2.16 | 0.0311 |
| 689166 | 2013 | 1.79 | 1688 | 2.06 | 0.13 | -2.05 | 0.04 |
| 689166 | 2018 | 1.8 | 1939 | 2.13 | 0.13 | -2.57 | 0.0103 |
| 689166 | 2023 | 1.8 | 3004 | 2.09 | 0.129 | -2.27 | 0.0235 |
| 743174 | 2004 | 1.9 | 1174 | 2.11 | 0.0965 | -2.19 | 0.0288 |
| 743174 | 2014 | 1.93 | 1580 | 2.15 | 0.104 | -2.1 | 0.0357 |
| 749166 | 2008 | 1.81 | 1708 | 2.13 | 0.145 | -2.22 | 0.0264 |
| 749166 | 2013 | 1.88 | 1852 | 2.12 | 0.0965 | -2.51 | 0.0121 |
| 749166 | 2018 | 1.97 | 3290 | 2.19 | 0.107 | -2.11 | 0.0352 |
| 749166 | 2023 | 1.89 | 4715 | 2.18 | 0.132 | -2.2 | 0.0277 |
| 767158 | 2008 | 1.95 | 1639 | 2.2 | 0.111 | -2.29 | 0.0218 |
| 779142 | 2007 | 1.77 | 2597 | 2.19 | 0.192 | -2.2 | 0.0278 |
| 779142 | 2016 | 1.74 | 1173 | 2.03 | 0.136 | -2.11 | 0.0347 |
| 803142 | 2013 | 1.82 | 2307 | 2.23 | 0.149 | -2.78 | 0.00549 |

**Table S2**. Linear model output per site for the studied period 2003 - 2023. Table shows 27 negative and 10 positive slopes corresponding to 73% and 27% of the significant trends (8.2% = 37 out of 450 sites had significant trend).

| No | id_site | Slope | *P-value* | Trend direction |
| --- | --- | --- | --- | --- |
| 1 | 509150 | -0.0144 | 0.0479 | Decreasing |
| 2 | 533182 | -0.0101 | 0.0493 | Decreasing |
| 3 | 533198 | -0.017 | 0.0371 | Decreasing |
| 4 | 557166 | -0.0237 | 0.0379 | Decreasing |
| 5 | 569222 | -0.0208 | 0.0394 | Decreasing |
| 6 | 587238 | -0.00896 | 0.0086 | Decreasing |
| 7 | 599206 | -0.0112 | 0.000259 | Decreasing |
| 8 | 605150 | -0.019 | 0.0234 | Decreasing |
| 9 | 605198 | -0.0332 | 0.0109 | Decreasing |
| 10 | 611238 | -0.00494 | 0.0367 | Decreasing |
| 11 | 617134 | -0.00841 | 0.0312 | Decreasing |
| 12 | 635190 | -0.0101 | 0.0288 | Decreasing |
| 13 | 659206 | -0.0102 | 0.00354 | Decreasing |
| 14 | 659254 | -0.0423 | 0.0493 | Decreasing |
| 15 | 659270 | -0.00981 | 0.0461 | Decreasing |
| 16 | 671254 | -0.0164 | 0.000226 | Decreasing |
| 17 | 689214 | -0.0128 | 0.0231 | Decreasing |
| 18 | 701198 | -0.0105 | 0.0136 | Decreasing |
| 19 | 701262 | -0.0187 | 0.0366 | Decreasing |
| 20 | 701278 | -0.0211 | 0.00774 | Decreasing |
| 21 | 725278 | -0.00679 | 0.0021 | Decreasing |
| 22 | 731158 | -0.0215 | 0.0103 | Decreasing |
| 23 | 731270 | -0.0132 | 0.0157 | Decreasing |
| 24 | 743190 | -0.00309 | 0.0446 | Decreasing |
| 25 | 749182 | -0.0276 | 0.0376 | Decreasing |
| 26 | 749214 | -0.0178 | 0.019 | Decreasing |
| 27 | 773182 | -0.0106 | 0.0328 | Decreasing |
| 28 | 503142 | 0.0116 | 0.0398 | Increasing |
| 29 | 599126 | 0.00736 | 0.000165 | Increasing |
| 30 | 647254 | 0.0145 | 0.0425 | Increasing |
| 31 | 695174 | 0.0244 | 0.0273 | Increasing |
| 32 | 725118 | 0.00513 | 0.0113 | Increasing |
| 33 | 725182 | 0.0148 | 0.023 | Increasing |
| 34 | 725198 | 0.0049 | 0.0383 | Increasing |
| 35 | 737182 | 0.0069 | 0.0133 | Increasing |
| 36 | 767190 | 0.00791 | 0.00939 | Increasing |
| 37 | 827190 | 0.0197 | 0.00893 | Increasing |

**Table S3**. Linear mixed-effects model fit by REML. Response variable is Community Weighted Length Index (CWLI).

| Response variable | Explanatory variables | Estimate | Std.Error | *df* | t value | *p value* |
| --- | --- | --- | --- | --- | --- | --- |
| CWLI | (Intercept) | 11.941 | 1.556 | 1419 | 7.675 | <0.001 |
|  | year | -0.005 | 0.001 | 1419 | -6.147 | <0.001 |
|  | elevation | -0.006 | 0.001 | 448 | -5.184 | <0.001 |
|  | year*elevation | 0.000003 | 0.000 | 1419 | 5.081 | <0.001 |

**Table S4**. Linear model with abundance as response variable.

| Response variable | Explanatory variables | Estimate | Std.Error | z value | Pr(>\|z\|) |
| --- | --- | --- | --- | --- | --- |
| Abundance | (Intercept) | 0.67 | 0.003104 | 214.85 | <0.001 |
|  | elevation | 0.06 | 0.003064 | 18.25 | <0.001 |
|  | wing length | -0.04 | 0.003126 | -12.308 | <0.001 |
|  | elevation*wing length | -0.03 | 0.003161 | -9.784 | <0.001 |

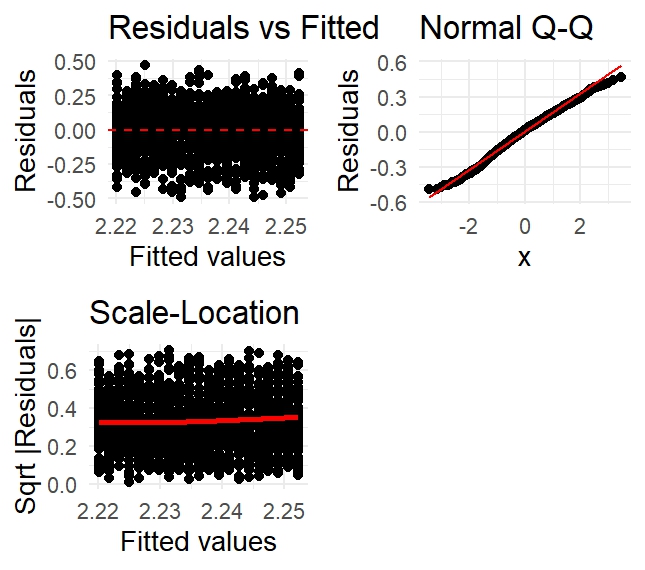

**Figure S1. The diagnostic plots assess the validity of the linear regression assumptions for Community Weighted Length Index (CWLI) trends. The *Residuals vs Fitted* plot checks for heteroscedasticity, where a random scatter indicates constant variance. The *Normal Q-Q Plot* evaluates residual normality, with points aligning to the reference line suggesting normally distributed errors. The *Scale-Location Plot* further tests variance homogeneity, where a flat trend supports homoscedasticity.**

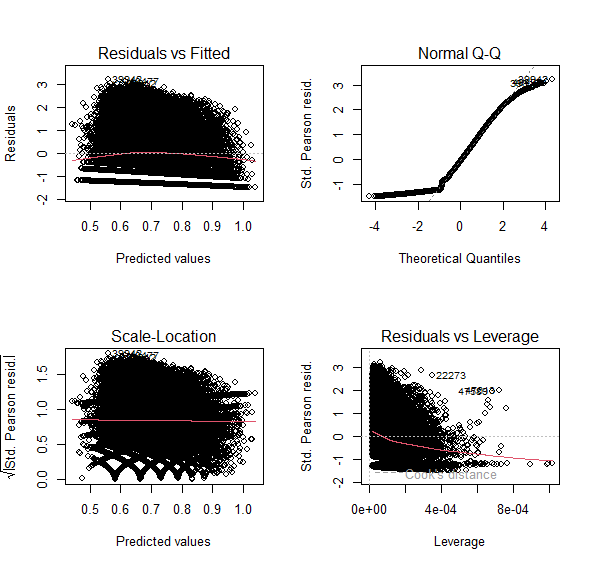

**Figure S2. The diagnostic plots assess the assumptions of the linear mixed-effects model analyzing Community Weighted Length Index (CWLI) trends across sites. The *Residuals vs Fitted Plot* checks for homoscedasticity, where a random scatter of points suggests consistent variance. The *Normal Q-Q Plot* evaluates the normality of residuals, with points closely following the reference line indicating normally distributed errors. The *Scale-Location Plot* further tests variance homogeneity, where a flat trend supports homoscedasticity and the *Residuals vs Leverage Plot* identifies influential data points with Cook’s Distance to help detect these points.**

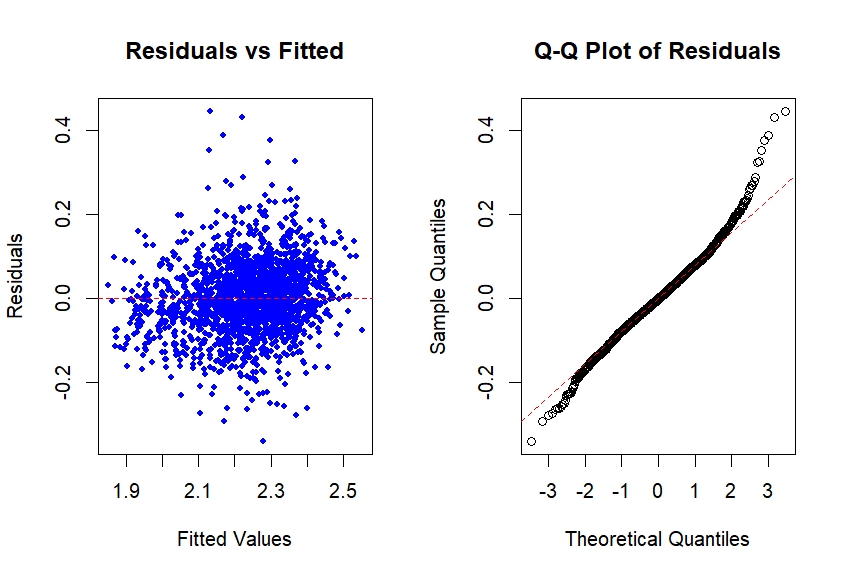

**Figure S3. The diagnostic plots assess the assumptions of the negative binomial model analyzing abundance trends across elevation and time. The *Residuals vs Fitted Plot* checks for homoscedasticity, where a random scatter of points suggests consistent variance. The *Normal Q-Q Plot* evaluates the normality of residuals, with points closely following the reference line indicating normally distributed errors.**
